# A dendritic guidance receptor functions in both ligand dependent and independent modes

**DOI:** 10.1101/2025.02.11.637350

**Authors:** Anay R Reddy, Sebastian J Machera, Zoe T Cook, Engin Özkan, Kang Shen

**Affiliations:** Department of Biology, Stanford University, Stanford, CA 94305, USA; Howard Hughes Medical Institute, Stanford University, Stanford, CA 94305, USA; Neurosciences IDP, Stanford University, Stanford, CA 94305, USA; Department of Biochemistry and Molecular Biology, The University of Chicago, Chicago, IL 60637, USA; Institute for Neuroscience, The University of Chicago, Chicago, IL 60637, USA; Institute for Biophysical Dynamics, The University of Chicago, Chicago, IL 60637, USA

## Abstract

The formation of an appropriately shaped dendritic arbor is critical for a neuron to receive information. Dendritic morphogenesis is a dynamic process involving growth, branching, and retraction. How the growth and stabilization of dendrites are coordinated at the molecular level remains a key question in developmental neurobiology. The highly arborized and stereotyped dendritic arbors of the *Caenorhabditis elegans* PVD neuron are shaped by the transmembrane DMA-1 receptor through its interaction with a tripartite ligand complex consisting of SAX-7, MNR-1, and LECT-2. However, receptor null mutants exhibit strongly reduced dendrite outgrowth, whereas ligand null mutants show disordered branch patterns, suggesting a ligand-independent function of the receptor. To test this idea, we identified point mutations in *dma-1* that disrupt receptor-ligand binding and introduced corresponding mutations into the endogenous gene. We show that the ligand-free receptor is sufficient to drive robust, disordered dendritic branch formation but results in a complete loss of arbor shape. This disordered outgrowth program utilizes similar downstream effectors as the stereotyped outgrowth program, further arguing that ligand binding is not necessary for outgrowth. Finally, we demonstrate that ligand binding is required to maintain higher-order dendrites after development is complete. Taken together, our findings support a surprising model in which ligand-free and ligand-bound DMA-1 receptors have distinct functions: the ligand-free receptor promotes stochastic outgrowth and branching, whereas the ligand-bound receptor guides stereotyped dendrite morphology by stabilizing arbors at target locations.

## Introduction

A neuron’s ability to receive synaptic inputs or sensory stimuli is dependent on dendritic morphology. Size, shape, and branching pattern are key parameters of dendrite arbors that vary widely across neuronal cell types. During development, dendrite morphogenesis requires the integration of extrinsic signals derived from the nearby complex tissue environment, which ultimately instructs dendrite shape, with intrinsic cytoskeletal rearrangements to drive dendrite outgrowth (Valnegri and Puram et al., 2015). A critical question in developmental neurobiology is how extrinsic and intrinsic mechanisms converge to form and maintain stereotyped dendritic arbors.

Guidance cues serve as extrinsic signals to instruct the growth direction of major dendritic processes during dendrite morphogenesis in a manner that resembles axon development. For example, a putative gradient of the Sema3A ligand near the marginal zone attracts apical dendrites of hippocampal CA1 pyramidal neurons towards the pial surface in mice (Polleux et al., 2000). Similarly, high levels of the Slit ligand at the central nervous system midline in *Drosophila melanogaster* repels motoneuron dendrites (Mauss et al., 2009). The Sema3A and Slit ligands exert their instructive roles on dendrite arborization through binding their cognate receptors neuropilin-1 and Robo, respectively (Polleux et al., 2000; Furrer and Kim et al., 2003). Similar findings studying other ligand-receptor systems, such as Netrin/Frazzled and Wnt/Frizzled, supported a model in which cognate guidance receptors are activated by ligand binding, thereby triggering downstream intracellular effectors to regulate dendrite outgrowth (Valnegri and Puram et al., 2015). These examples suggest that diffusible gradients of attractants and repellants guide the morphology of major dendritic branches.

Additionally, fine dendritic branch morphologies must be continuously refined to establish a non-overlapping dendritic field between the same class of neurons. Therefore, these neurons employ self-avoidance and dendritic tiling mechanisms to minimize overlap (Grueber and Sagasti, 2010). These growth repulsive mechanisms ensure maximum coverage of a target field while minimizing overlap between branches. Repulsion between branches is achieved through *cis* and *trans* homophilic interactions between alternatively-spliced cell adhesion molecules, including DSCAM and protocadherins (Millard and Zipursky, 2008; Kostadinov and Sanes, 2015). While the mechanisms regulating spacing between fine dendritic branches are well-understood, the molecular mechanisms that drive pathfinding of fine dendritic branches remain unclear. Furthermore, how receptors regulate the cytoskeletal machinery to refine the shape of fine branches is not fully understood.

Guidance receptors regulate dendrite outgrowth through the recruitment and activation of cytoskeletal regulators. For example, receptor-mediated dendrite outgrowth can employ the Rho GTPases RhoA, Rac1, and Cdc42 (Luo, 2002). These molecules cycle between an inactive GDP-bound state and an active GTP-bound state (Etienne-Manneville and Hall, 2002). In the GTP-bound active state, Rho GTPases promote actin nucleation and polymerization to drive outgrowth (Jan and Jan 2010; Etienne-Manneville and Hall, 2002). Transitioning from the inactive conformation to the active state requires guanine nucleotide exchange factors (GEFs) to facilitate the replacement of GDP with GTP (Etienne-Manneville and Hall, 2002). Guidance receptors play a crucial role in promoting GTPase activity by recruiting GEFs. For example, upon binding to an ephrinB ligand, the EphB2 receptor phosphorylates and recruits the Rac GEF Tiam1, which promotes dendritic spine development (Tolias et al., 2007). Therefore, receptors can bridge extrinsic cues with intrinsic outgrowth programs by recruiting cytoskeletal regulators. Whether intrinsic outgrowth programs can be activated through a receptor but independently of extrinsic signals remains unclear.

To further understand the relationship between ligand-receptor interaction and dendrite development, we generated a ligand-free receptor to test whether dendrite outgrowth of a highly organized arbor can occur in the absence of ligand-receptor binding. We employed an *in silico* approach to design a putative non-binding mutant receptor and validated that ligand-receptor binding is abrogated through a biochemical approach. Genetic experiments reveal that the non-binding receptor is sufficient to drive disordered dendrite formation that phenocopies ligand null conditions. The disordered outgrowth phenotypes in non-binding receptor mutants require the same downstream effectors utilized in stereotyped outgrowth of organized arbors, arguing that ligand binding is not required for receptor-mediated activation of cytoskeletal regulators. Finally, we demonstrate that ligand-receptor binding is particularly required for stabilization and maintenance of higher-order branches. Taken together, these results argue that ligand binding of receptor is dispensable for outgrowth but is required for stabilizing dendrites at appropriate locations.

## Results

### PVD dendrite arborization requires both extrinsic cues and intrinsic outgrowth programs

The PVD mechanosensory neuron in the nematode *Caenorhabditis elegans* (*C. elegans*) affords a powerful experimental system to study dendrite morphogenesis and dendrite branch patterning. PVD has an expansive dendritic arbor with orthogonally oriented primary, secondary, tertiary, and quaternary dendrites (Albeg et al., 2011) The higher-order secondary, tertiary, and quaternary branches form structures resembling menorahs (Albeg et al., 2011). Menorah formation requires three extrinsic cues: the cell adhesion molecules SAX-7/L1CAM and MNR-1/Menorin expressed by the hypodermis, and the chemotaxin LECT-2/LECT2 secreted from the muscle cells (Zou et al., 2018; Salzberg et al., 2013; Dong et al., 2013; Zou et al., 2016). All three ligands collectively form a tripartite ligand complex and bind to the extracellular domain of the DMA-1 receptor expressed by PVD (Zou et al., 2016). The intracellular domain of DMA-1 promotes dendrite outgrowth by recruiting the Rac GEF TIAM-1/Tiam1 (Zou et al., 2018). DMA-1 also physically interacts with the claudin-like co-receptor HPO-30, which recruits the pentameric Wave Regulatory Complex (WRC) (Smith et al., 2013; Zou et al., 2018) through its cytoplasmic tail. Together, TIAM-1 activated Rac relieves self-inhibition of the WRC, which subsequently triggers Arp2/3 mediated actin assembly to drive dendrite outgrowth (Ding et al., 2022). These findings are consistent with a model in which DMA-1 bridges extrinsic and intrinsic programs by promoting cytoskeletal changes in a manner dependent upon ligand binding (***Figure 1B***). In this model, the tripartite ligand complex localizes to specific tissues to guide a deterministic outgrowth program which leads to a stereotyped dendritic arbor. In support of this model, both ligand null and receptor null mutants completely lack menorahs. However, ligand null mutants display robust but disordered branches, whereas receptor null mutants have sparse higher-order branching, arguing that the receptor can trigger dendrite outgrowth independently of ligand binding (Zou et al., 2016; Liu et al., 2011). Furthermore, time-lapse microscopy of developing PVD neurons reveals that some secondary dendrite outgrowth events are stochastic in direction and can retract back to the primary dendrite. These findings demonstrate that PVD dendrite development is not fully deterministic and challenge the canonical ligand-receptor model in PVD dendrite morphogenesis (Smith et al., 2010).

**Figure 1.**
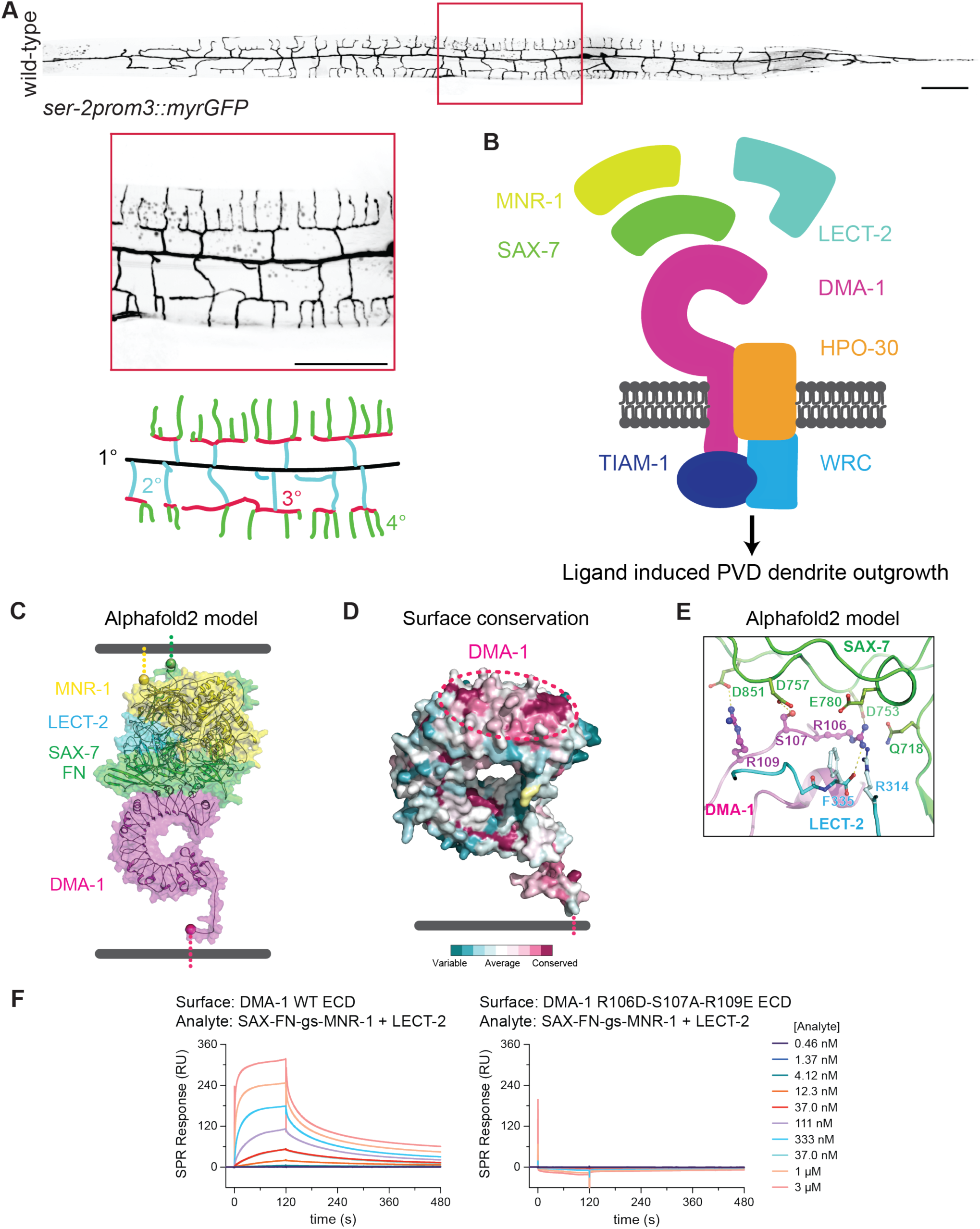
Three receptor residues are necessary for DMA-1 receptor binding to a tripartite ligand complex. (**A**) (Top) Lateral fluorescence z-projections of the entire PVD neuron visualized with a *ser-2prom3::myristoylated::GFP* membrane marker in a young adult animal. Red box indicates region 150 μm anterior of PVD cell body. Scale bar, 100μm. (Middle) Inset enlargement of boxed region. Scale bar, 50 μm. (Bottom) Schematic of PVD dendrite branching. Primary (1°) dendrite shown in black, secondary (2°) dendrites shown in blue, tertiary (3°) dendrites shown in red, quaternary (4°) dendrites shown in green. (**B**) Original model depicting ligand-receptor binding bridging extrinsic signals with intrinsic cytoskeletal changes to activate PVD dendrite outgrowth. In this model, DMA-1 receptor binds an extracellular tripartite ligand complex consisting of SAX-7, LECT-2, and MNR-1, which triggers activation of actin polymerization through the downstream Rac GEF TIAM-1, coreceptor HPO-30, and Wave Regulatory Complex (WRC). (**C**) Alphafold 2 (multimer) model of DMA-1 ectodomain + LECT-2 + SAX-7 FN domains + MNR-1. The Alphafold2 model orients the C-terminal ends of MNR-1 and SAX-7 at the expected site for the skin cell membrane. (**D**) Surface conservation of DMA-1 highlights the ligand engagement surface as highly conserved (dashed oval). (**E**) Close-up of residues DMA-1 R106 to R109 in the Alphafold 2 model. (**F**) SPR sensorgrams for the binding of the tripartite ligand complex on wild-type and mutant DMA-1 ectodomain (ECD).

### Three DMA-1 residues are critical for ligand-receptor binding

Previous work has shown that deletion of the extracellular Leucine Rich Region (LRR) domain of DMA-1, which mediates ligand binding, caused robust but disorganized dendrite branching (Shi, Ho, and Tao et al., 2024). This finding argued that the ligand complex’s interaction with DMA-1 is required for dendrite patterning but is dispensable for dendrite outgrowth. To test this idea more precisely, we sought to generate point mutations in the extracellular domain of DMA-1 to abolish ligand-receptor binding. Since there is no published structure of the ligand-receptor complex, we leveraged Alphafold and surface conservation to predict the structure of the ligand-receptor complex and identify candidate residues on the receptor required for ligand binding (Jumper, Evans, Pritzel et al., 2021). The Alphafold2 model identified the N-terminal convex side of the DMA-1 LRR domain as the ligand-interaction surface (***Figure 1C***). Furthermore, surface conservation analysis showed the same region as the most conserved surface on DMA-1 (***Figure 1D***). Analysis of the interface in the predicted models identified that the DMA-1 residues R106, S107, and R109 were likely critical for binding to the SAX-7 ligand (***Figure 1E***).

To create and validate a non-ligand interacting DMA-1 mutant *in vitro*, we established large-scale expression systems for the DMA-1 ectodomain, LECT-2, and a single-chain version of SAX-7 Fibronectin type III (FN) domains linked with MNR-1, as MNR-1 could not be expressed alone in significant quantities. Using purified proteins and surface plasmon resonance (SPR), we showed that wild-type DMA-1 interacts strongly with a 1:1 SAX-7 FN-MNR-1:LECT-2 mixture (***Figure 1F***). When we engineered and tested the DMA-1 R106D-S107A-R109E mutant with SPR, we observed no ligand binding, validating that these residues are required for DMA-1 to bind the ligand complex (***Figure 1F***).

### DMA-1 can promote disorganized outgrowth independently of ligand binding

We used clustered regularly interspersed short palindromic repeats (CRISPR)-Cas9 genome editing to generate a non-binding receptor mutant allele (Jinek, Chylinski et al., 2012, Dickinson et al. 2016). We first asked how ligand-free receptor affects dendrite patterning and outgrowth. Given that ligand null mutants exhibit disordered dendrite outgrowth whereas receptor null mutants display little dendrite outgrowth, we predicted that PVD morphology in *dma-1* (*non-binding*) mutants would be distinct from *dma-1* null animals while resembling *sax-7* null animals. The *ser-2prom3::myristoylatedGFP* marker was used to label the PVD neuron, and young adult animals were imaged (Liu et al., 2011; ***Figure 1A***). We first compared the dendrite morphologies of *dma-1* (*non-binding*) with those of *dma-1* (*null*) and *sax-7* (*null*). All three mutant classes were similar in that they failed to form quaternary branches when compared to wild-type animals (***Figure 2A-D*** and ***2I*).** We next quantified higher-order dendrite length in the proximal anterior dendrite. Interestingly, the dendrite outgrowth of *dma-1* (*non-binding*) animals phenocopied *sax-7* (*null*) but were distinct from *dma-1* (*null*) animals, which display severely reduced outgrowth (***Figure 2B-D*** and ***2J*).** Furthermore, *dma-1* (*non-binding*); *sax-7* (*null*) double mutants showed similar outgrowth phenotypes to the corresponding single mutants, arguing that our engineered point mutations indeed disrupt SAX-7 binding (***Figure 2D-E*** and ***2J*).** Taken together, these data demonstrate that SAX-7 ligand binding to DMA-1 is required for dendrite patterning but is dispensable for dendrite outgrowth.

**Figure 2.**
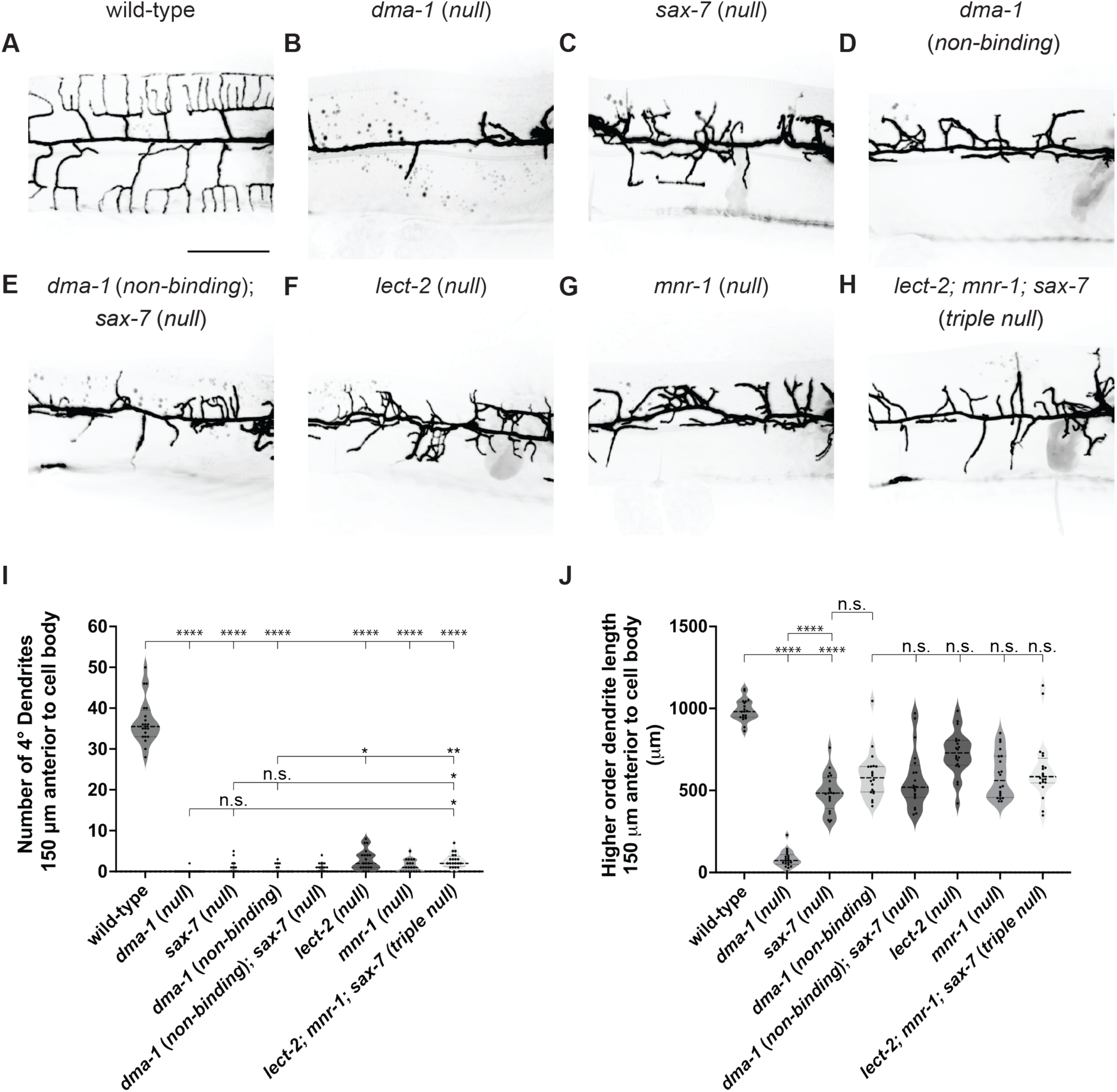
Ligand-free DMA-1 receptor can promote disordered dendrite arborization. (**A-H**) Lateral fluorescence z-projections of PVD neuron 150 μm anterior to the PVD cell body in young adult animals of the genotypes indicated. PVD labelled with a *ser-2prom3::myristoylated::GFP* membrane marker. Scale bar, 50 μm. (**I-J**) Quantifications of number of quaternary (4°) dendrites (**I**) and total higher-order dendrite length (**J**) 150 μm anterior to the PVD cell body. Medians are represented in thick dashed lines and quartiles are represented in thin dashed lines. P values were calculated using a Brown-Forsythe and Welch one-way ANOVA with Dunnett’s test. n=20 for all conditions. n.s., (not significant), p>0.05, *p≤0.05, **p≤0.01, ****p≤0.0001.

Given that SAX-7 is one component of the tripartite ligand complex, we next asked whether the loss of either the LECT-2 or MNR-1 ligands would resemble the disruption of SAX-7 binding to DMA-1 in terms of disorganized dendritic outgrowth. We compared both higher-order dendrite patterning and dendrite length in *dma-1* (*non-binding*), *lect-2* (*null*), *mnr-1* (*null*), and *lect-2; mnr-1; sax-7* triple null animals (***Figure 2D, F-H***). All single mutant animals displayed a similar inability to form ordered quaternary branches when compared to wild-type, with *lect-2* (*null*) showing slightly increased quaternary branches when compared to *dma-1* (*non-binding*). While the *lect-2; mnr-1; sax-7* triple null animals also exhibited reduced quaternary branching when compared to wild-type, these animals displayed a slightly increased number of quaternaries when compared to the *sax-7* (*null*) and *dma-1* (*non-binding*) single mutant classes (***Figure 2I***). All mutant classes exhibited similar disordered dendrite outgrowth phenotypes in that they showed less total dendrite outgrowth than wild-type but more outgrowth than *dma-1* (*null*) (***Figure 2J***). Taken together, these data support a model in which ligand-free receptor can promote disorganized dendritic outgrowth and ligand-bound receptor is required for appropriate dendritic morphology.

To directly understand how the SAX-7 interaction with DMA-1 instructs PVD morphology, we endogenously tagged the SAX-7 ligand with mNeonGreen to determine the extent of dendrite misalignment with ligand in wild-type and *dma-1* (*non-binding*) animals. SAX-7 is particularly enriched along ventral and dorsal sublateral tracts which coincide with where PVD tertiary dendrites form. Therefore, misalignment was scored by measuring the length of SAX-7 tracts along the ventral and dorsal sublateral lines that do not colocalize with dendrites (***Figure 3A, D***). While the dendrites of wild-type animals mostly grew along the sublateral lines, leading to a low misalignment score, the misalignment in *dma-1* (*non-binding*) animals was far greater (***Figure 3B, C, E*).** Interestingly, we observed that some secondary dendrites in wild-type animals did not overlap with SAX-7 tracts (***Figure 3C***, white arrows). In contrast to secondaries that overlapped with SAX-7 tracts, secondaries that deviated from ligand appeared to exhibit an atypical and non-perpendicular angle relative to the primary dendrite. These results strengthen our claim that the engineered point mutations disrupt SAX-7-DMA-1 interaction and that SAX-7 binding to DMA-1 is not necessary to stimulate dendrite outgrowth. Furthermore, the reduced colocalization of SAX-7 sublateral tracts with PVD dendrites in mutants supports the idea that SAX-7 ultimately instructs the dendrite shape through its interaction with DMA-1. Finally, these data demonstrate that dendrites can grow without contacting SAX-7 tracts, albeit in a less stereotyped manner.

**Figure 3.**
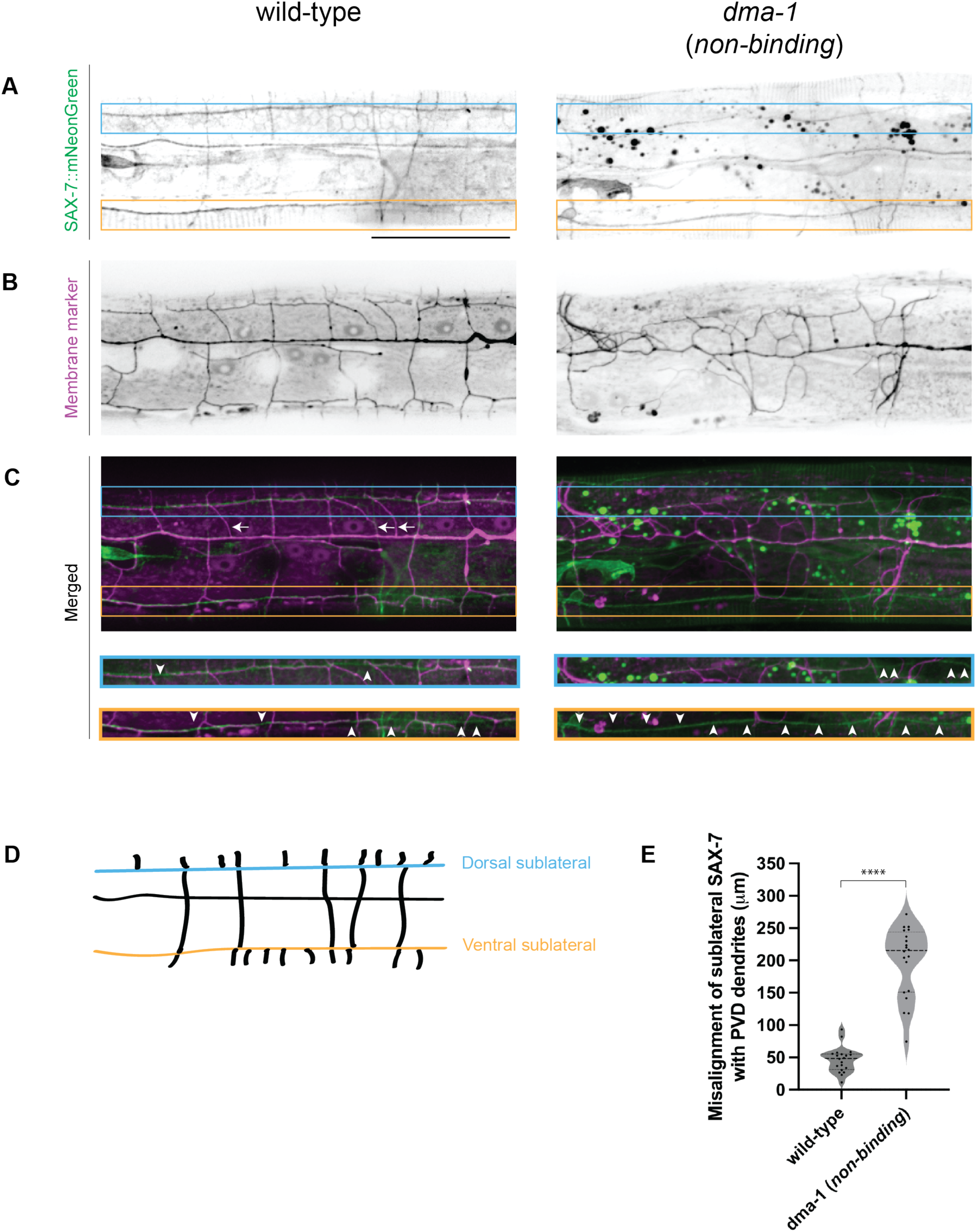
Ligand-free DMA-1 receptor fails to align PVD dendrites with SAX-7 ligand tracts. (**A-C**) Lateral fluorescence z-projections of endogenously labelled SAX-7 (**A**) and PVD morphology (**B**) 150 μm anterior to PVD cell body in young adult animals. PVD labelled with a *ser-2prom3::myristoylated::mCherry* membrane marker. Merged images shown in (**C**). Left: wild-type representative image. Right: *dma-1* (*non-binding*) representative image. Blue and orange boxes and insets denote dorsal and ventral sublateral lines, respectively. White arrows denote wild-type secondary dendrites that partially or completely fail to align with SAX-7 tracts. White arrowheads denote regions at the sublateral lines where PVD dendrites fail to align with SAX-7 tracts. Scale bar, 50 μm. (**D**) Schematic of SAX-7 ligand tracts 150 μm anterior to PVD cell body. Blue and orange lines represent dorsal and ventral sublateral tracts, respectively. (**E**) Quantifications measuring length of SAX-7 ligand tract misalignment with PVD dendrites 150 μm anterior to PVD cell body. Medians are represented in thick dashed lines and quartiles are represented in thin dashed lines P value was calculated using a two-tailed unpaired *t*-test with Welch’s correction. n=20 for both conditions. ****, p≤0.0001.

### Ligand-free and ligand-bound DMA-1 engage similar downstream effectors

Previous work identified that DMA-1 physically interacts with the coreceptor HPO-30, which is also required for PVD dendrite morphogenesis (Zou et al., 2018). The cytosolic tail of DMA-1 and HPO-30 directly bind to the RAC guanine nucleotide exchange factor (GEF) TIAM-1 and the WAVE regulatory complex (WRC), respectively. Together, TIAM-1 and WRC promote actin polymerization by activating the Arp2/3 complex (Kramer et al., 2023). During the formation of quaternary dendrites, the DMA-1/HPO-30 complex organizes F-actin to promote dendrite outgrowth and branching (Shi et al., 2021). Based on our work arguing that ligand-receptor binding is not required for dendrite outgrowth, we next asked whether ligand-free receptor also functioned through HPO-30 and TIAM-1 or if outgrowth was achieved through different interactors.

In a *dma-1* (*non-binding*) strain, we introduced null mutations in either *hpo-30* or *tiam-1* to determine if disordered dendrite outgrowth is dependent on the function of these genes. All mutant classes exhibited reduced dendrite outgrowth when compared to wild-type animals (***Figure 4A,G***). Importantly, the *dma-1* (*non-binding*); *hpo-30* (*null*) and *dma-1* (*non-binding*); *tiam-1* (*null*) double mutants both showed diminished dendrite outgrowth when compared to *dma-1* (*non-*binding) single mutants (***Figure 4D-G***) or the *hpo-30* (*null*) and *tiam-1* (*null*) single mutant classes (***Figure 4B-C, G***). These data demonstrate that ligand-free receptor promotes dendrite outgrowth through the HPO-30 coreceptor and TIAM-1 Rac GEF. Furthermore, our results support the surprising notion that ligand-receptor binding is not required to trigger downstream effectors that promote dendrite outgrowth and challenges the classic view that ligand-receptor signaling alone triggers neurite outgrowth.

**Figure 4.**
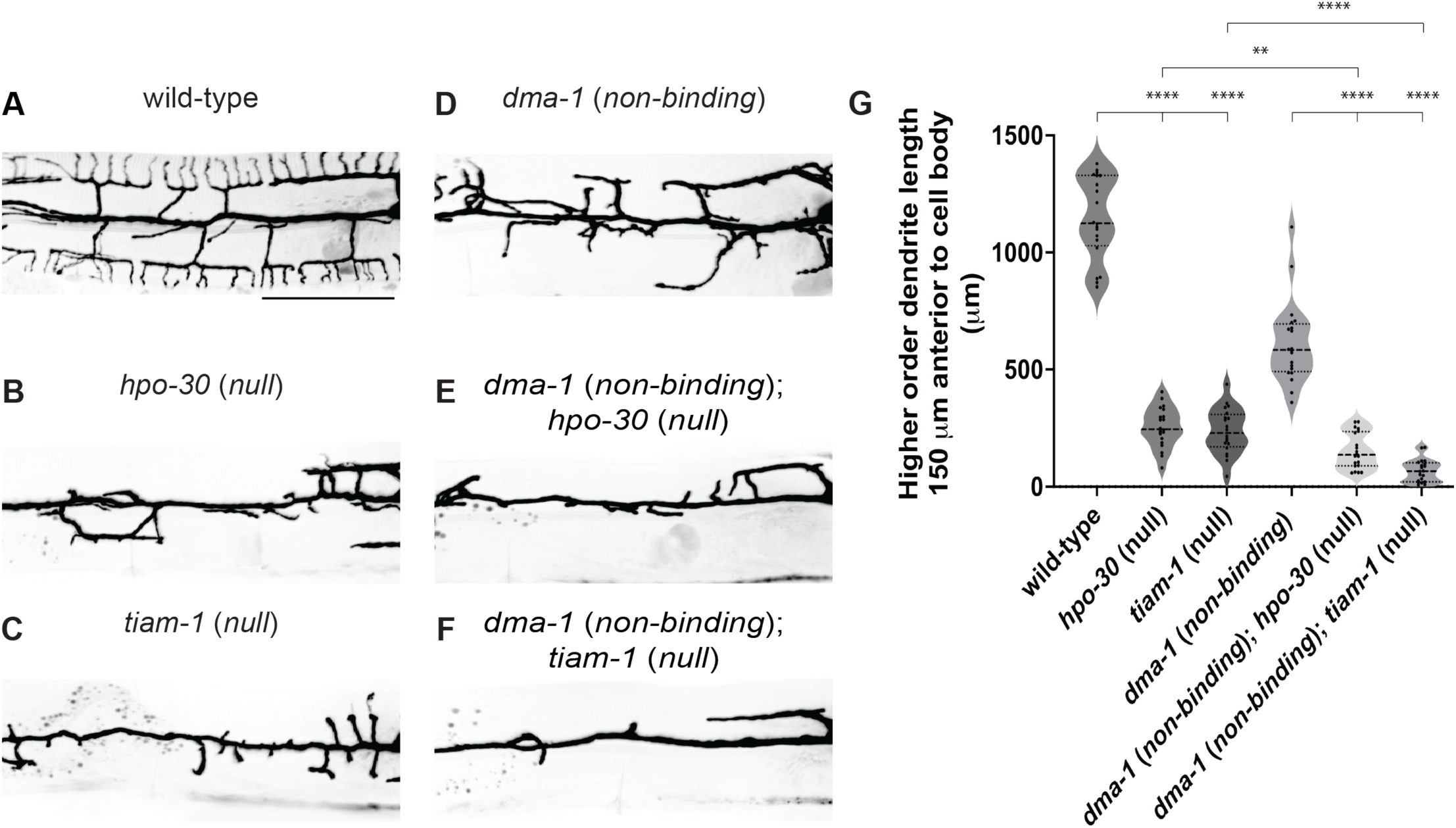
Ligand-free DMA-1 receptor promotes dendrite outgrowth through the Rac GEF TIAM-1 and coreceptor HPO-30. (**A-F**) Lateral fluorescence z-projections of PVD neuron 150 μm anterior to the PVD cell body in young adult animals of the genotypes indicated. PVD labelled with a *ser-2prom3::myristoylated::GFP* membrane marker. Scale bar, 50 μm. (**G**) Quantifications of total higher-order dendrite branching length 150 μm anterior to the PVD cell body. Medians are represented in thick dashed lines and quartiles are represented in thin dashed lines. P values were calculated using a Brown-Forsythe and Welch one-way ANOVA with Dunnett’s test. n=20 for all conditions. **p≤0.01, ****p≤0.0001.

### Ligand binding is essential for dendrite stability and maintenance

Given that ligand-free receptor can promote dendrite outgrowth through canonical downstream effectors, we next asked how ligand-bound receptor uniquely regulates PVD morphology. Since the higher-order tertiary and quaternary branches are reduced in ligand null and *dma-1* (*non-binding*) mutant classes, we speculated that ligand-receptor binding is specifically required to maintain the stereotyped menorah shape (Salzberg et al., 2013; Dong et al., 2013, ***Figure 2B, D, I***). We formulated a model in which the ligand binding state of DMA-1 confers distinct roles: ligand-free receptors exclusively promote outgrowth, whereas ligand-bound receptors exclusively promote PVD patterning by stabilizing higher-order branches along ligand tracts.

To first test whether DMA-1 has role in maintaining PVD higher-order dendrites after outgrowth, we used a temperature sensitive allele of *dma-1* to temporally downregulate the receptor after menorah formation. The *dma-1* (*ts*) allele harbors a point mutation in the extracellular LRR domain (S324L). Animals cultured at the permissive temperature (16°) from hatching to adulthood display menorah formation, albeit with reduced quaternaries when compared to wild-type animals (***Figure 5B, D***). To test if DMA-1 is required for dendrite stabilization, we cultured both wild-type and *dma-1* (*ts*) animals at the permissive temperature. Upon reaching the young adult stage, when menorah formation is complete, animals were either kept at the permissive temperature or transferred to the restrictive temperature (25°). After 24 hours, morphologies of Day 1 adults were scored (***Figure 5A***). While the shift to the restrictive temperature had no effect on dendrite morphology in wild-type animals, *dma-1* (*ts*) animals exhibited a reduced number of tertiary and quaternary branches (***Figure 5B-D***). These data demonstrate that DMA-1 is not only required for PVD outgrowth but also for dendrite branch maintenance.

**Figure 5.**
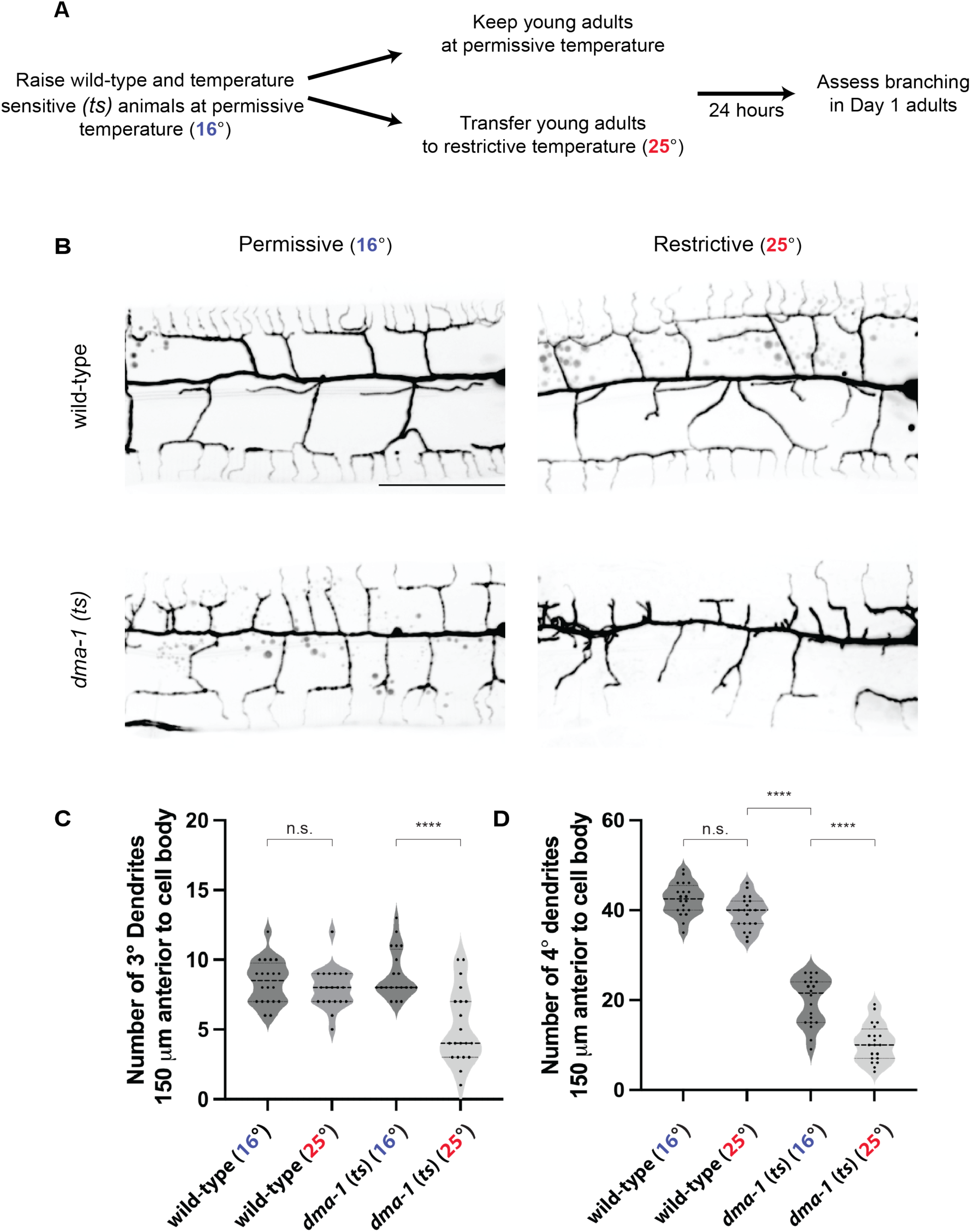
The DMA-1 receptor is required for higher-order dendrite branch maintenance. (**A**) Experimental schematic to determine how temporal downregulation of DMA-1 receptor after PVD outgrowth affects higher-order dendrite branching. (**B**) Lateral fluorescence z-projections of PVD neuron 150 μm anterior to the PVD cell body in Day 1 adult animals. PVD labelled with a *ser-2prom3::myristoylated::GFP* membrane marker. Top, representative wild-type animals. Bottom, representative *dma-1* temperature sensitive (*ts*) animals. Left, animals cultured at permissive temperature (16°). Right, animals transferred to restrictive temperature (25°) at the young adult stage. Scale bar, 50 μm. (**C-D**) Quantifications of number of tertiary (3°) dendrites (**C**) and quaternary (4°) dendrites (**D**) 150 μm anterior to the PVD cell body. Medians are represented in thick dashed lines and quartiles are represented in thin dashed lines. P values were calculated using a Brown-Forsythe and Welch one-way ANOVA with Dunnett’s test. n=20 for all conditions. n.s., (not significant), p>0.05, ****p≤0.0001.

We hypothesized that only ligand-bound DMA-1 receptor could promote branch stabilization. To test this model, we generated a *dma-1* allele where an auxin inducible degron (AID*) sequence was inserted into an endogenously tagged *dma-1::GFP* allele (Eichel et al., 2022). In tandem with a pan-somatic transgene expressing the F-box protein TIR1, our degron tagged *dma-1::GFP::AID\** allele conferred the ability to temporally degrade receptor upon addition of the plant hormone auxin (Ashley et al., 2021). We determined that 17 hours of auxin treatment was sufficient to induce significant degradation of DMA-1 based on loss of the endogenous GFP signal (***Supplementary Figure 1***). To assess if the induced degradation affected DMA-1 function, we treated young adult *dma-1::GFP::AID\** animals with 10 mM auxin and observed a loss of quaternary branches (***Figure 6B’, C***). We next tested whether ligand-free receptor could promote higher-order branch stabilization. Given that *dma-1* (*non-binding*) homozygous animals are unable to form higher-order branches, we generated transheterozygotes that had one copy of a non-degradable *dma-1* (*non-binding*) allele and one copy of the *dma-1::GFP::AID\** allele (***Figure 2I*, *Figure 6A***). These transheterozygotes provided a system in which higher-order branches could form by the young adult stage, allowing us to test the role of ligand binding in dendrite maintenance (***Figure 6B’’’***). As a control, we generated a second class of transheterozygotes containing one copy of a wild-type, non-degradable *dma-1* allele and one copy of the *dma-1::GFP::AID\** allele (***Figure 6A***). We predicted that only the transheterozygotes with a non-degradable wild-type allele would maintain dendrite branches after the depletion of degradable receptor. Temporal downregulation of the AID* tagged pool of receptor, which can bind ligand, was induced by transferring young adult transheterozygotes from auxin free plates to 10 mM auxin plates, and Day 1 adults were imaged and scored for quaternary branches (***Figure 6A***). With this experimental paradigm, the majority of remaining receptor in the transheterozygotes would either be wild-type or *dma-1* (*non-binding*) after degradation. Transheterozygotes expressing a non-degradable wild-type receptor allele exhibited no loss of quaternary branching after auxin treatment (***Figure 6B’’, C***). In contrast, transheterozygotes expressing the non-binding receptor allele showed reduced quaternary branches after auxin treatment (***Figure 6B’’’, C***). Taken together, these data argue that ligand-receptor binding is required for the stabilization and maintenance of higher-order dendrites.

**Figure 6.**
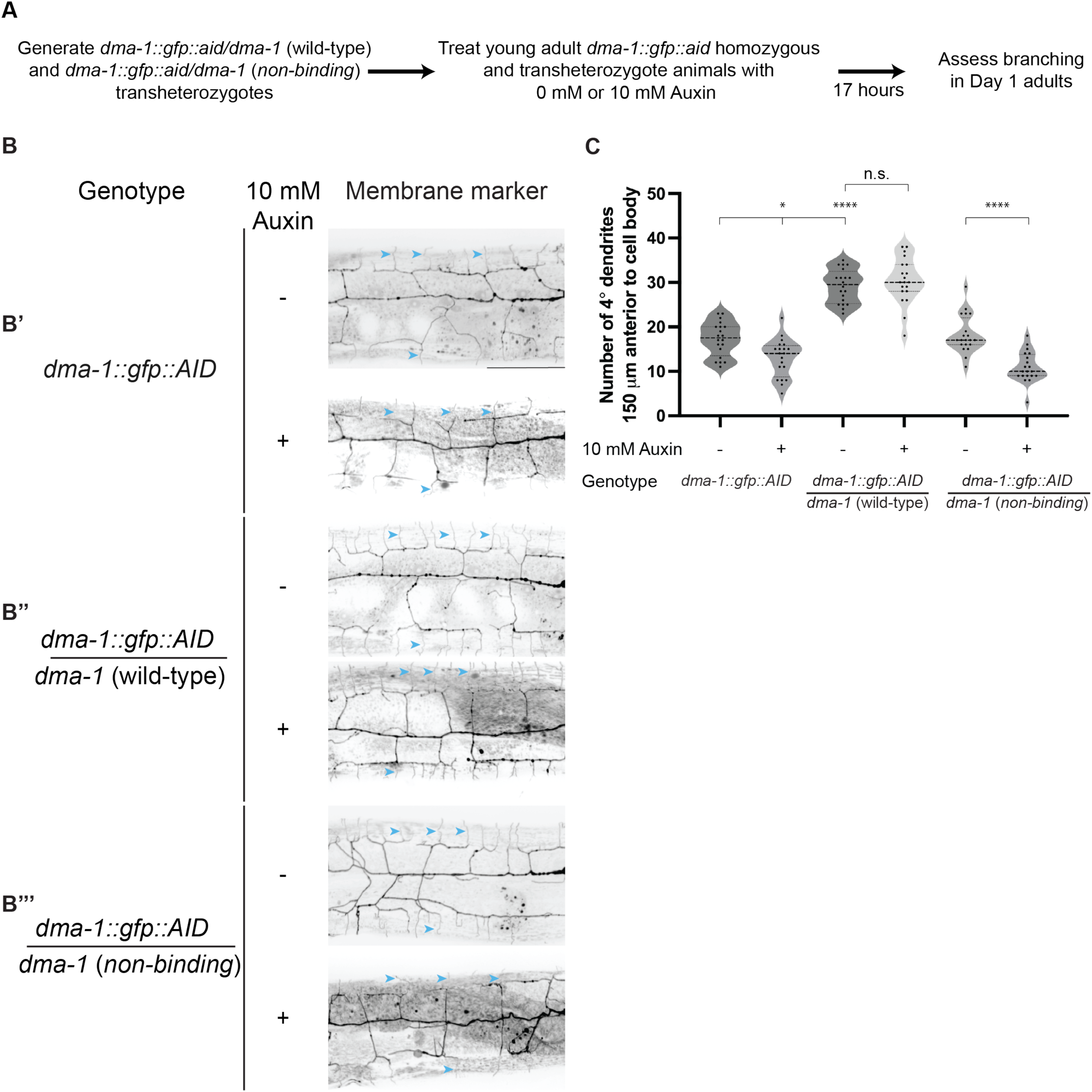
The DMA-1 receptor promotes higher-order branch stabilization through ligand binding. (**A**) Experimental schematic to determine whether ligand-free receptor promotes branch stabilization. (**B**) Lateral fluorescence maximum intensity z-projections of PVD in Day 1 adult animals. PVD labelled with a *ser-2prom3::myristoylated::mCherry* membrane marker. B’, *dma-1::gfp::aid* homozygous animals. B’’, *dma-1::gfp::aid*/*dma-1* (wild-type) transheterozygotes. B’’’, *dma-1::gfp::aid*/*dma-1* (*non-binding*) transheterozygotes. Top images in B’, B’’, and B’’’ are representative images of animals that did not receive auxin treatment. Bottom images are representative images of animals that received 10 mM auxin treatment at the young adult stage. Blue arrowheads denote examples of quaternary branches. Scale bar, 50 μm. (**C**) Quantifications of quaternary (4°) branches 150 μm anterior to the PVD cell body. Medians are represented in thick dashed lines and quartiles are represented in thin dashed lines. P values were calculated using a Brown-Forsythe and Welch one-way ANOVA with Dunnett’s test. n=20 for all conditions. n.s., (not significant), p>0.05, *p≤0.05,**** p≤0.0001.

## Discussion

The acquisition of appropriate dendritic morphology is critical for a neuron’s ability to receive information. Underlying this process is the coordination between growth and stabilization of dendritic branches during development. A critical question in developmental neurobiology is how both dendrite growth and stabilization are regulated by guidance molecules and their cognate receptors. One possibility is that molecular guidance cues promote stereotyped outgrowth by binding to a receptor to direct dendrites to target locations. Alternatively, molecular guidance cues may function stochastically by binding to a receptor to selectively stabilize dendrite branches. The stereotyped growth model predicts that guidance receptor activation leads to directed outgrowth, and that the loss of guidance cues severely inhibits dendrite outgrowth. In contrast, the selective stabilization model predicts that only a fraction of the dendrite outgrowth events leads to a stabilized branch, and that the loss of guidance cues will prevent stabilization of branches at target locations while having either a small or no effect on dendrite outgrowth (Shree and Sutradhar et al., 2022; Palavalli et al., 2021). Here, we propose that the PVD neuron employs both stereotyped growth and selective stabilization programs in the formation of its dendritic arbor. Our findings reveal that the DMA-1 receptor can regulate both these programs: in the ligand-free state, DMA-1 promotes stochastic outgrowth; in the ligand-bound state, DMA-1 promotes stereotyped dendritic morphology by stabilizing dendrites along ligand tracts, which enables further dendritic outgrowth at these regions (***Figure 7***).

**Figure 7.**
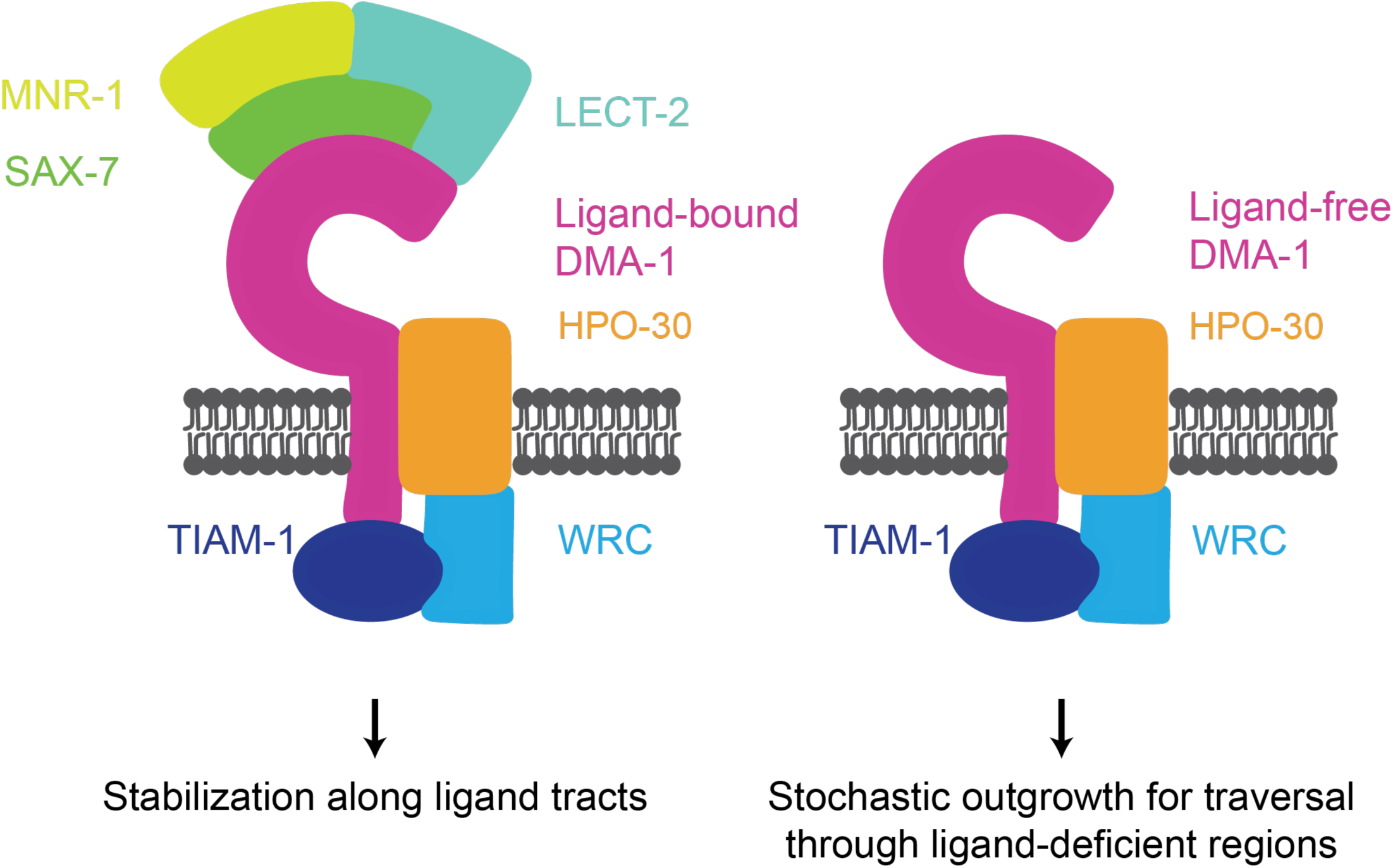
Model for how ligand-bound and ligand-free receptor pools promote PVD dendrite arborization. DMA-1 receptor binding to the SAX-7/LECT-2/MNR-1 tripartite ligand complex promotes both stereotyped dendrite outgrowth along ligand tracts and branch stabilization. Ligand-free receptor drives stochastic dendrite outgrowth branching through ligand deficient regions. Both programs function through the downstream effectors TIAM-1 and HPO-30.

Our claim that a receptor can promote dendritic outgrowth without binding ligand is supported by several lines of evidence. Although ligand null mutants lack the menorah structure indicative of ordered growth, they can establish a disorganized dendritic arbor that likely arises from stochastic dendritic outgrowth (***Figure 2I, J***). Furthermore, mutants in which the extracellular LRR domain of DMA-1 was deleted also exhibited disorganized branching (Shi, Ho, and Tao et al., 2024). In this work, we demonstrate that the substitution of three residues in the LRR domain is sufficient to disrupt ligand binding, and that the disorganized dendritic morphologies by mutants expressing this non-binding receptor allele phenocopies those found in ligand null mutants (***Figure 1E-F*, *Figure 2***). Examining ligand localization lends further support for the claim that ligand-free receptors can promote dendrite outgrowth. The SAX-7 ligand is distributed unevenly along the epidermal cells upon which PVD elaborates its dendrites: SAX-7 is low in regions where secondary dendrites extend but is high in regions where tertiary and quaternary dendrites develop (Dong et al., 2013). These findings are inconsistent with the model that dendrite outgrowth requires DMA-1 binding to its cognate ligands. Instead, we posit that some secondaries require ligand-free DMA-1 to drive stochastic outgrowth and require ligand-bound DMA-1 to stabilize at the sublateral lines.

If PVD employs both stereotyped growth and selective stabilization programs, how are these growth strategies coordinated? We propose that the availability of an extrinsic instructive factor dictates the balance between these programs. The development of the PVD primary dendrite is regulated by SAX-7 from the pioneering ALA axon, which interacts with both DMA-1 and SAX-3/Robo on PVD to promote both primary dendrite extension and fasciculation, respectively (Ramirez-Suarez et al., 2019; Chen et al., 2019). The apposition of externally derived SAX-7 with where the PVD primary dendrite extends and fasciculates argues that this initial mode of development mainly utilizes stereotyped growth programs. Unlike primary dendrite development, some secondary dendrites must traverse regions lacking SAX-7 ligand (***Figure 3C***, white arrows). In combination with other work, our findings support a model in which ligand-free DMA-1 can drive branch initiation and extension of these secondaries. In this model, secondary branches that cannot reach the sublateral line, where SAX-7, MNR-1, and LECT-2 are present, fail to stabilize and fully retract to the primary branch (Dong et al., 2013, Salzberg et al., 2013; Zou et al., 2016; Smith et al., 2010). In contrast, secondaries that reach the sublateral line can stabilize upon ligand-receptor binding (Salzberg et al., 2013; Dong et al., 2013). Therefore, we propose that the apparent stochastic growth of secondary branches is a consequence of low ligand levels in the region of extension, which promotes outgrowth driven by ligand-free DMA-1. On the other hand, formation of tertiary and quaternary dendrites occurs in regions abundant with ligands, enabling these branches to immediately stabilize after growth events. The presence of the tripartite ligand complex is used as a guidepost to instruct tertiary and quaternary dendrite outgrowth (Salzberg et al., 2013; Dong et al., 2013; Zou et al., 2016). We demonstrate that in addition to promoting higher-order branch outgrowth, ligand-bound receptor can promote branch stabilization (***Figure 2I*** and ***Figure 6C***). Consistent with our model, the development of secondary dendrites appears to exhibit more growth and retraction events when compared to that of quaternary dendrites (Smith et al., 2010; Tao et al., 2022). Therefore, PVD dendrite arborization employs two distinct programs in a spatially dependent manner: a stereotyped growth program is employed in regions enriched with instructive factors, whereas a stochastic outgrowth program is employed in regions that are deficient in instructive factors.

Our work contributes to a growing body of evidence that challenges the previously held notion in which guidance cues induce their cognate receptors to trigger downstream signaling pathways regulating dendrite outgrowth (Shi, Ho, and Tao et al., 2024). Instead, we assert that ligand-free receptor can promote disordered dendrite arborization. Surprisingly, our genetic experiments argue that ligand-receptor binding is not necessarily required for recruitment and activation of downstream cytoskeletal regulators. Finally, we demonstrate that ligand-receptor binding not only functions to promote ordered dendrite arborization but also serves to maintain higher-order branches at appropriate locations after development. Taken together, our findings support a model in which dendrite arborization can be achieved through a combination of cell-extrinsic and cell-intrinsic mechanisms. We propose that ligand-bound receptor stabilizes dendrites along ligand tracts to subsequently enable further dendritic outgrowth at appropriate regions, and that ligand-free receptor promotes stochastic outgrowth by promoting traversal through ligand-deficient areas. The balancing of these programs enables the PVD neuron to navigate a complex tissue environment and ultimately innervate appropriate regions to form a complex dendritic arbor.

## Materials and Methods

### Protein Structure Modeling

We used Alphafold 2 (version 2.3) locally, as implemented in Colabfold version 1.5.2 (Jumper et al., 2021; Evans et al., 2021; Kim et al., 2024), and Alphafold 3, made available by Deepmind at https://alphafoldserver.com (Abramson et al., 2024). Alphafold 2 reported an iPTM value of 0.806 for the predicted complex, while Alphafold 3 reported a 0.69 value. For surface conservation analysis, we used the Consurf server, available at https://consurf.tau.ac.il (Ashkenazy et al., 2016).

### Protein Expression and Purification

We used the baculoviral expression system for protein production *in vitro*. In short, we created baculoviruses using co-transfection of Sf9 cells with linearized baculoviral DNA with plasmids containing the protein of interest in the pAcGP67A backbone, which carries a gp64 signal peptide for protein secretion. The amplified viral stocks were used to infect High Five cells (*Trichoplusia ni*) grown in serum-free ESF-921 (Expression Systems). The secreted proteins, in which LECT-2 was N-terminally hexahistidine tagged and the remaining proteins were C-terminally hexahistidine tagged, were purified using Ni-NTA metal affinity chromatography, followed by size-exclusion chromatography on Superdex 200 Increase 10/300 columns in HEPES-buffered saline (HBS), containing 10 mM HEPES pH 7.4 and 150 mM NaCl.

For SPR experiments we expressed and purified the following four constructs: wild-type and mutated DMA-1 ectodomain (residues 20 to 505) with a C-terminal Avi-tag and hexahistidine tag; LECT-2 (residues 23 to 335) with an N-terminal hexahistidine tag followed by a tobacco etch virus protease site; and a single-chain fusion of SAX-7 Fibronectin type III domains (SAX-7 FN) (residues 633 to 1207) followed by a 34-residue (GGGS)_8_GG linker, MNR-1 with the GPI-anchor sequence removed (residues 17 to 457) and a C-terminal hexahistidine tag. All constructs carried an N-terminal gp64 signal peptide for secretion.

### Surface Plasmon Resonance

For SPR, we used a Biacore T200 machine and a streptavidin-coupled Series S Sensor Chip SA (Cytiva). Wild-type and mutant DMA-1 ectodomains were biotinylated at the biotin acceptor peptide (i.e. Avi-tag) using BirA biotin ligase. Approximately 200 response units of wild-type and mutant DMA-1 were captured on separate channels, and 1:1 mixed SAX-7 FN–MNR-1:LECT-2 samples were run over both channels in a buffer containing 10 mM HEPES, pH 7.4, 150 mM NaCl and 0.05% Tween-20.

### *C. elegans* culture and strains

*C. elegans* were cultured at 20°C on NGM plates using OP50 *Escherichia coli* as a food source according to standard procedures unless otherwise noted (Stiernagle, 2006). Two constructs were used as PVD membrane markers: *ser2prom3::myristoylated::GFP* (*wyIs*592) and *ser2prom3::myristoylated::mCherry* (*wyIs*581). All strains and primers used in this study are listed in the Key resources table.

### Generation of genome-edited strains

Point mutations and endogenous fluorophore or degron insertions were created by gonadal microinjection of CRISPR-Cas9 protein complexes. For the generation of the *dma-1* (*non-binding*) allele, a single-stranded DNA primer consisting of 30 nucleotide homology arms was used as a repair template. This repair template included a silent site mutation conferring an *HaeIII* restriction enzyme cut site and removed the PAM site to prevent Cas9 from editing the repair template. SAX-7 was C-terminally tagged with mNeonGreen::AID*. The repair template for this insertion was generated by PCR amplification from the pJW2171 plasmid and contained a GSGGGG linker between the mNeonGreen and AID* sequences (Ashley et al., 2021). Generation of the *dma-1::GFP::AID\** allele was achieved by inserting an AID* sequence to the C terminus of the GFP sequence, which itself was previously inserted into the juxtamembrane region of DMA-1 (Eichel et al., 2022). The repair template for this insertion was generated by PCR amplification from the pJW2098 plasmid and included a GSGGGG linker upstream of the AID* sequence (Ashley et al., 2021).

CRISPR-Cas9 genome editing was performed using standard protocols (Dokshin et al., 2018). Cas9 protein and tracRNA (IDT) were both injected at 1.525 μM concentrations. sgRNAs (IDT) were injected at a concentration of 1.525 μM. Repair templates were injected at a concentration of 5 μM. To select for candidate animals with a successful genome edit, the pRF4 plasmid was injected at 50 ng μL^-1^. F_1_ roller animals were singled to fresh plates, and F_2_ animals were screened for the desired genome edit using PCR and Sanger sequencing. Primers used to amplify homology repair templates, sgRNA sequences, and genotyping primers used to verify generation of desired edits are listed in the Key resources table.

### *C. elegans* confocal microscopy and image analysis

Young and Day 1 adult hermaphrodite *C. elegans* animals were anaesthetized using 10 mM levamisole (Sigma-Aldrich) in M9 buffer and mounted on 3% agarose pads. With the exception of animals heat shocked at a 25°C incubator, all imaged animals were maintained in a 20°C incubator.

*C. elegans* animals were imaged on imaging systems consisting of either (1) an inverted Zeiss Axio Observer Z1 microscope paired with a Yokogawa CSU-X1 spinning-disk unit, a Hamamatsu EM-CCD digital camera, 488 nm and 561 nm solid state lasers, and a C-Apochromat 40x 0.9 NA objective controlled by Metamorph (version 7.8.12.0) or (2) an inverted Zeiss Axio Observer Z1 microscope paired with a Yokogawa CSU-W1 spinning-disk unit, a Prime 95B Scientific CMOS camera, 488 nm and 561 nm solid state lasers, and a C-Apochromat 40x 0.9NA objective controlled by 3i Slidebook (v6). The length of z-stacks was determined by manually setting top and bottom slices. In all cases, imaging conditions such as laser power, exposure time, and step-size (0.33 μm) were identical for all genotypes and conditions across the experiment.

Image analysis was performed on unprocessed images using Fiji software (Schindelin et al., 2022). Maximum-intensity and sum-intensity projections were rotated, cropped, and straightened to generate display images, and brightness and contrast were adjusted in Fiji. PVD micrographs of individual animals were stitched and cropped to show the region 150 μm anterior to the cell body using Fiji (Preibisch et al., 2009). All images are oriented in the following manner: the anterior-posterior axis is left to right, and ventral-dorsal axis is bottom to top.

The Simple Neurite Tracer (SNT) plugin was used in Fiji to calculate higher-order dendrite length (Arshadi et al., 2021). Briefly, non-overlapping traces were drawn on PVD higher-order dendrites for individual animals and saved. SNT was used to measure the length of individual traces.

### Statistical analysis

Statistical analysis was performed in GraphPad Prism 10. Animals were selected for measurements based on developmental stage, orientation on slide, and health. Sample size refers to the number of *C. elegans* animals imaged. For statistical tests, single pairwise comparisons of genotypes or treatments were analyzed using either two-tailed unpaired Student’s *t*-tests or two-tailed unpaired *t*-test with Welch’s correction. Multiple comparisons were performed using one-way analysis of variance (ANOVA) followed by post hoc Dunnett’s test. Figure legends indicate sample size, statistical tests used, and *p* values. All graphs prepared in GraphPad Prism.

## Acknowledgements

We would like to thank the past and current members of the Shen lab who provided excellent input in the writing of this manuscript. In particular, we thank Callista Yee, Kelsie Eichel, Yue Sun, Dane Kawano, and Junhao Xu for their feedback. We thank the trainees, staff, and faculty of the Stanford Department of Biology who provided helpful scientific discussions, technical assistance, and support. We also would like to thank the Stanford Superworm community for their encouragement and feedback. Finally, we thank Bryan Kirsch for critical reading of the manuscript.

## Key resources table

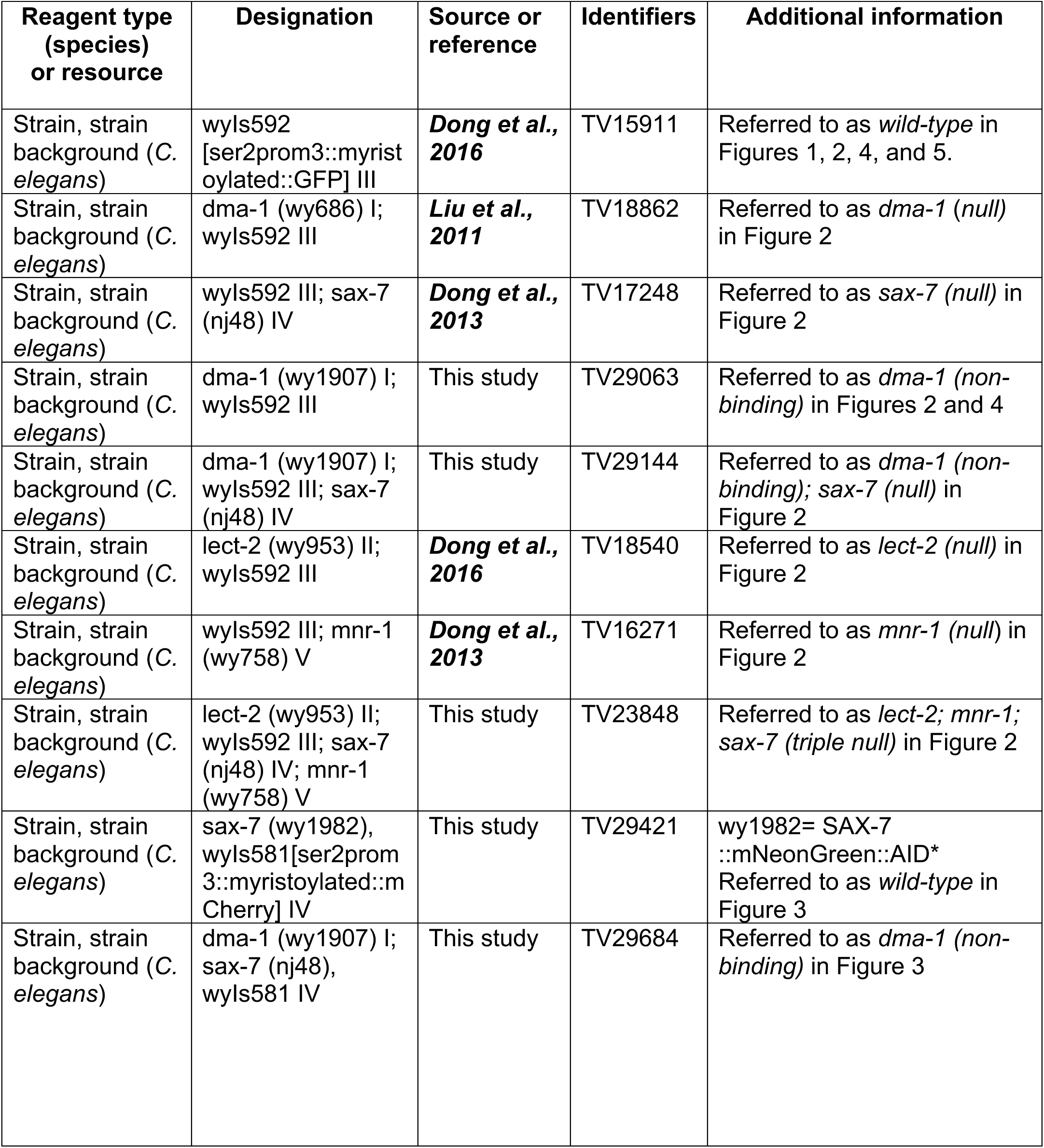
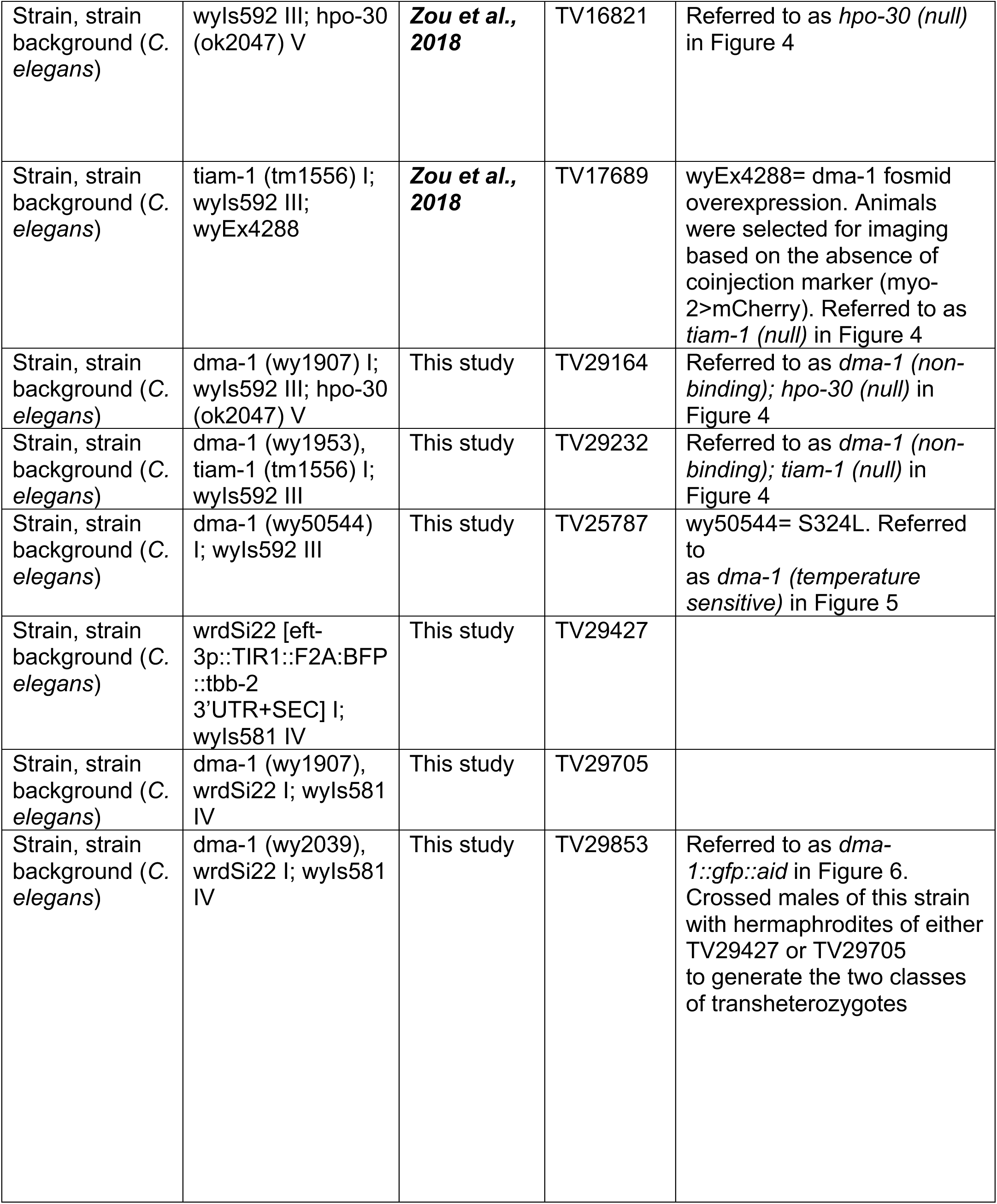
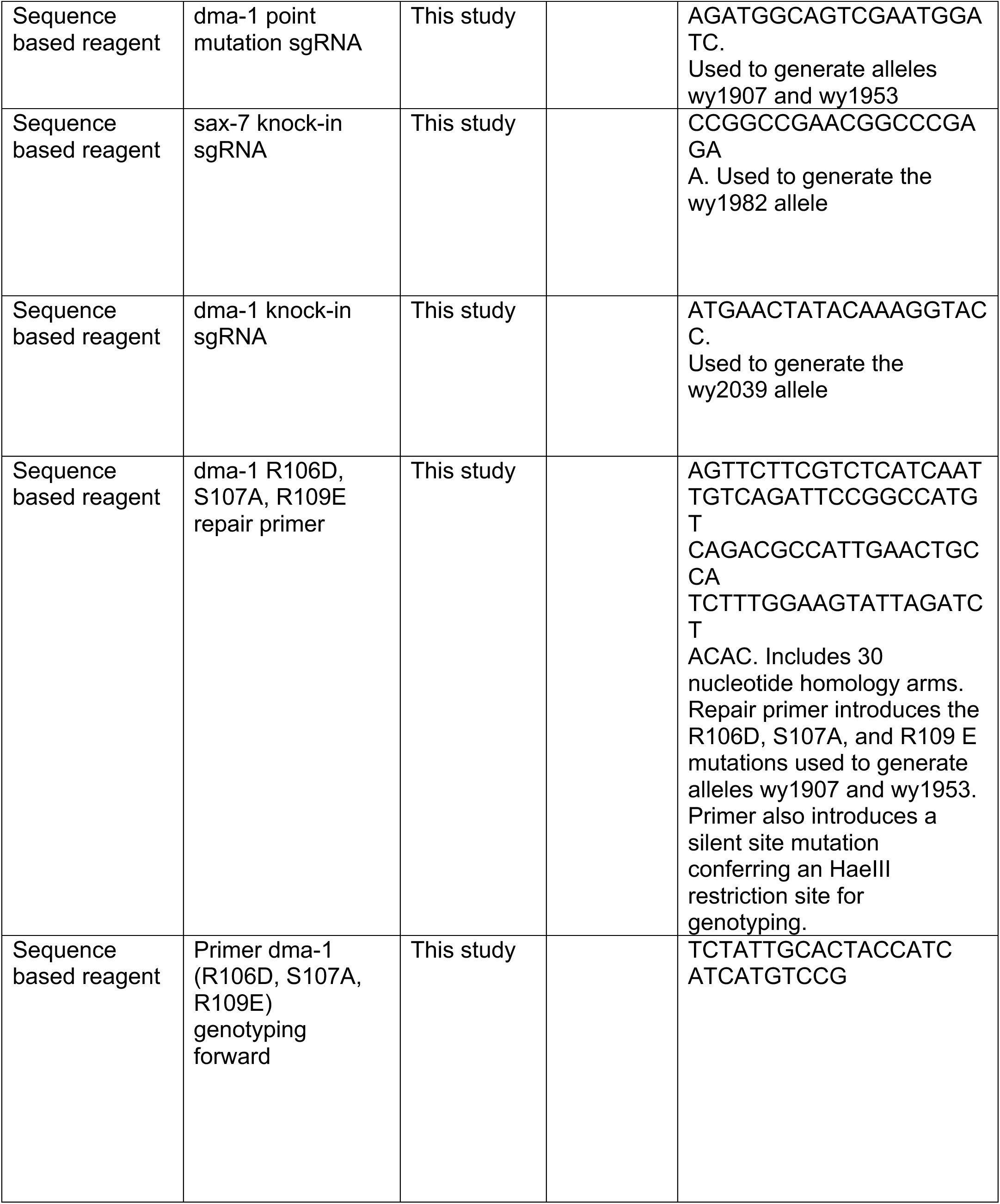
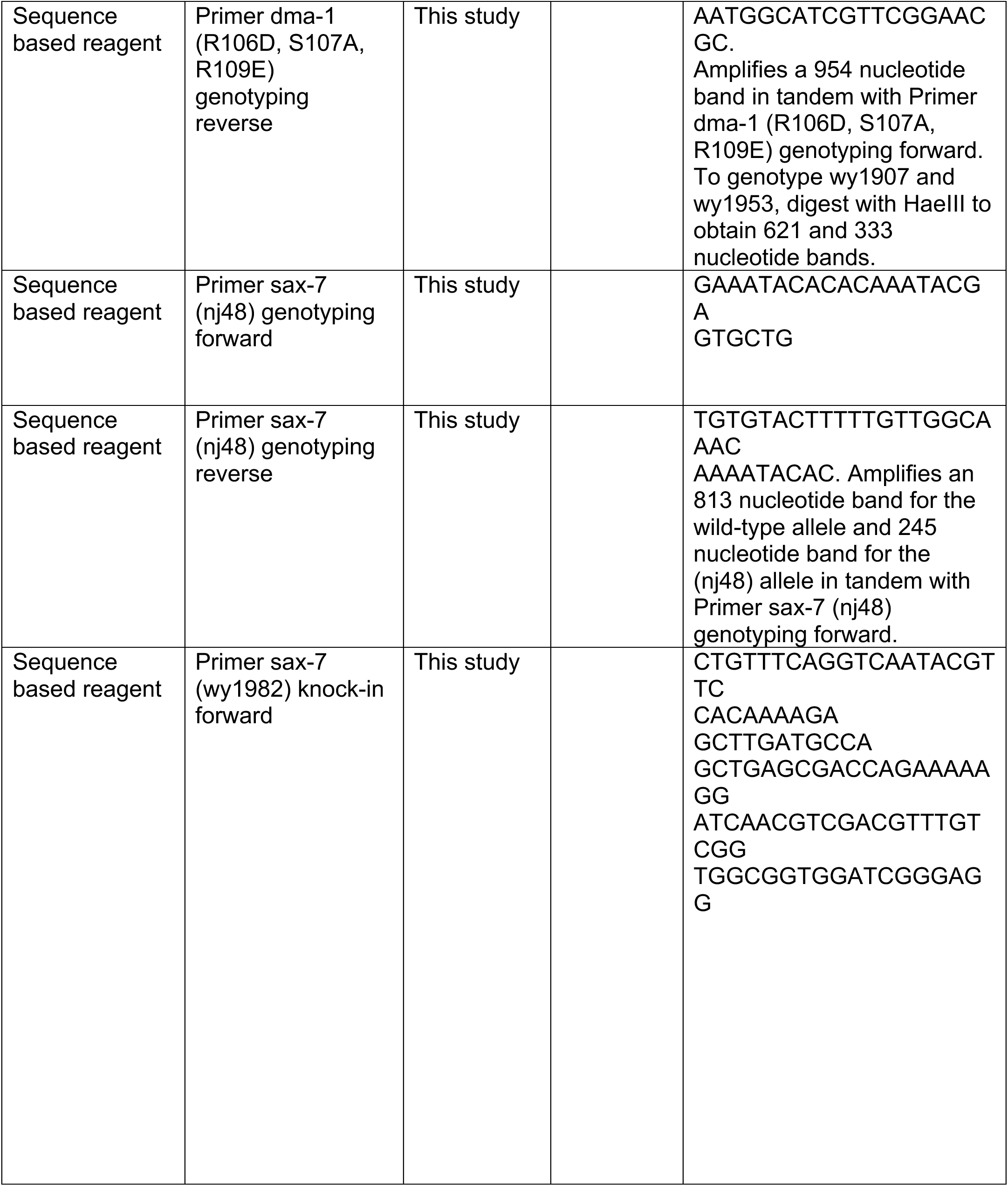
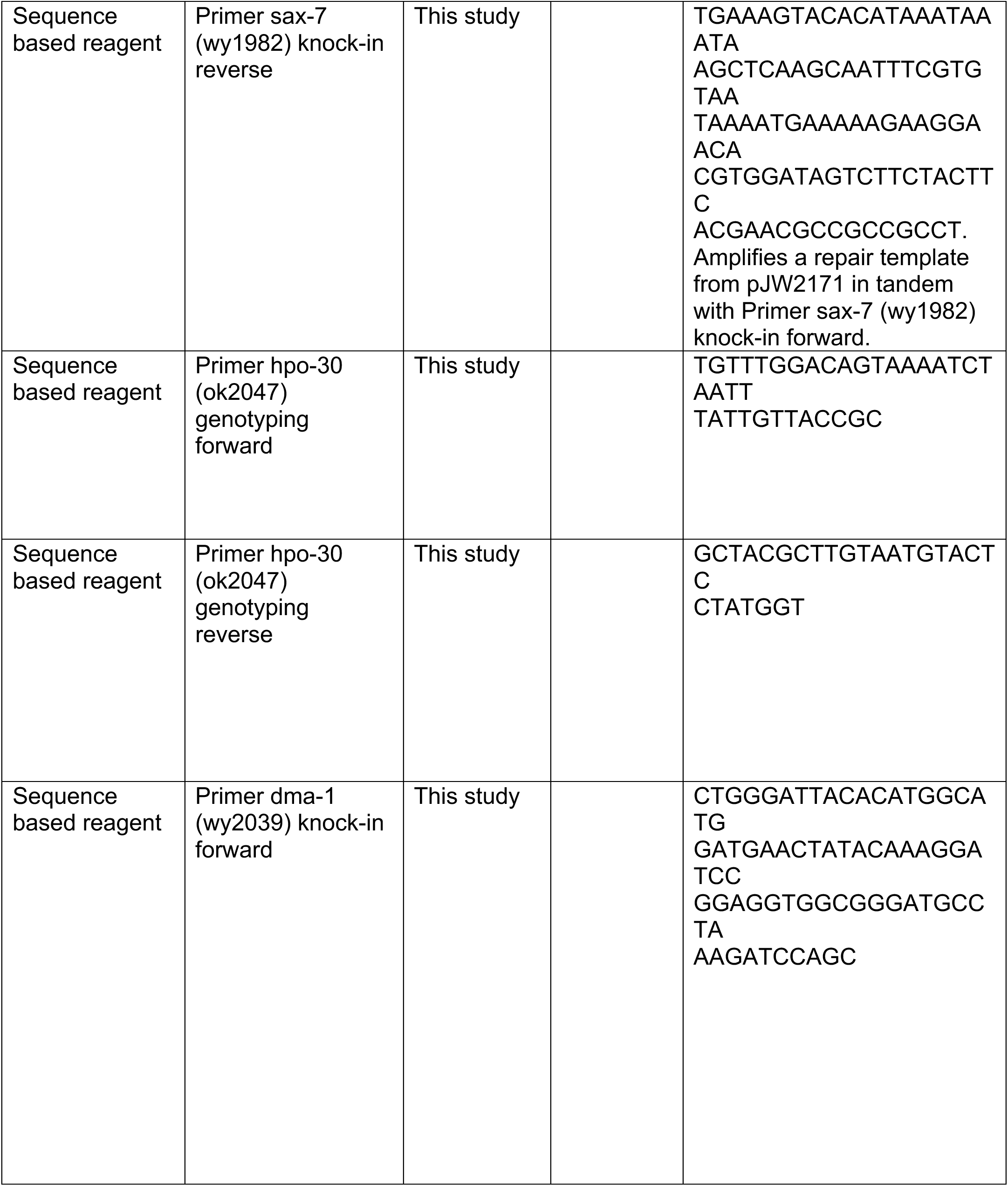
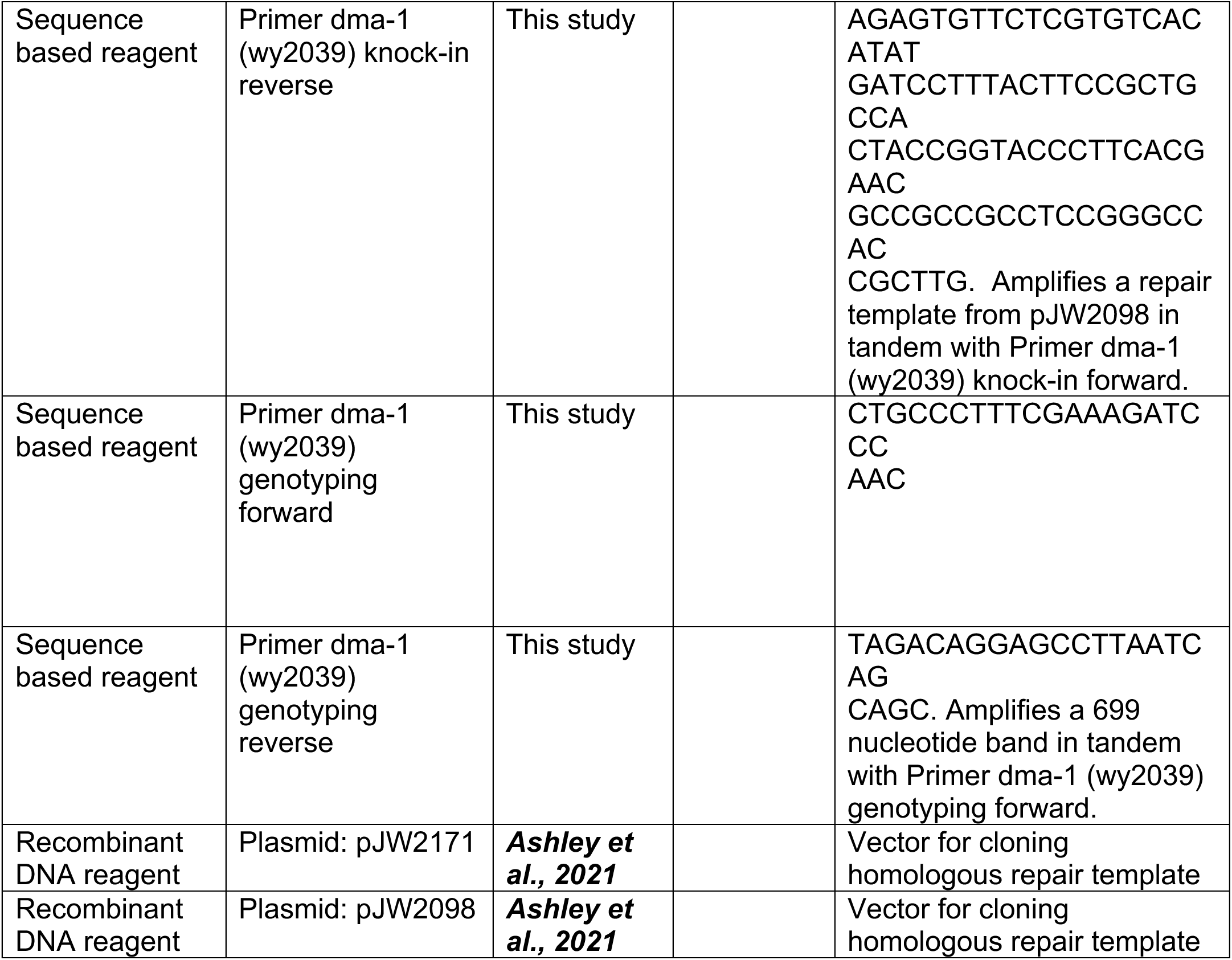

---

**Supplementary Figure 1.**
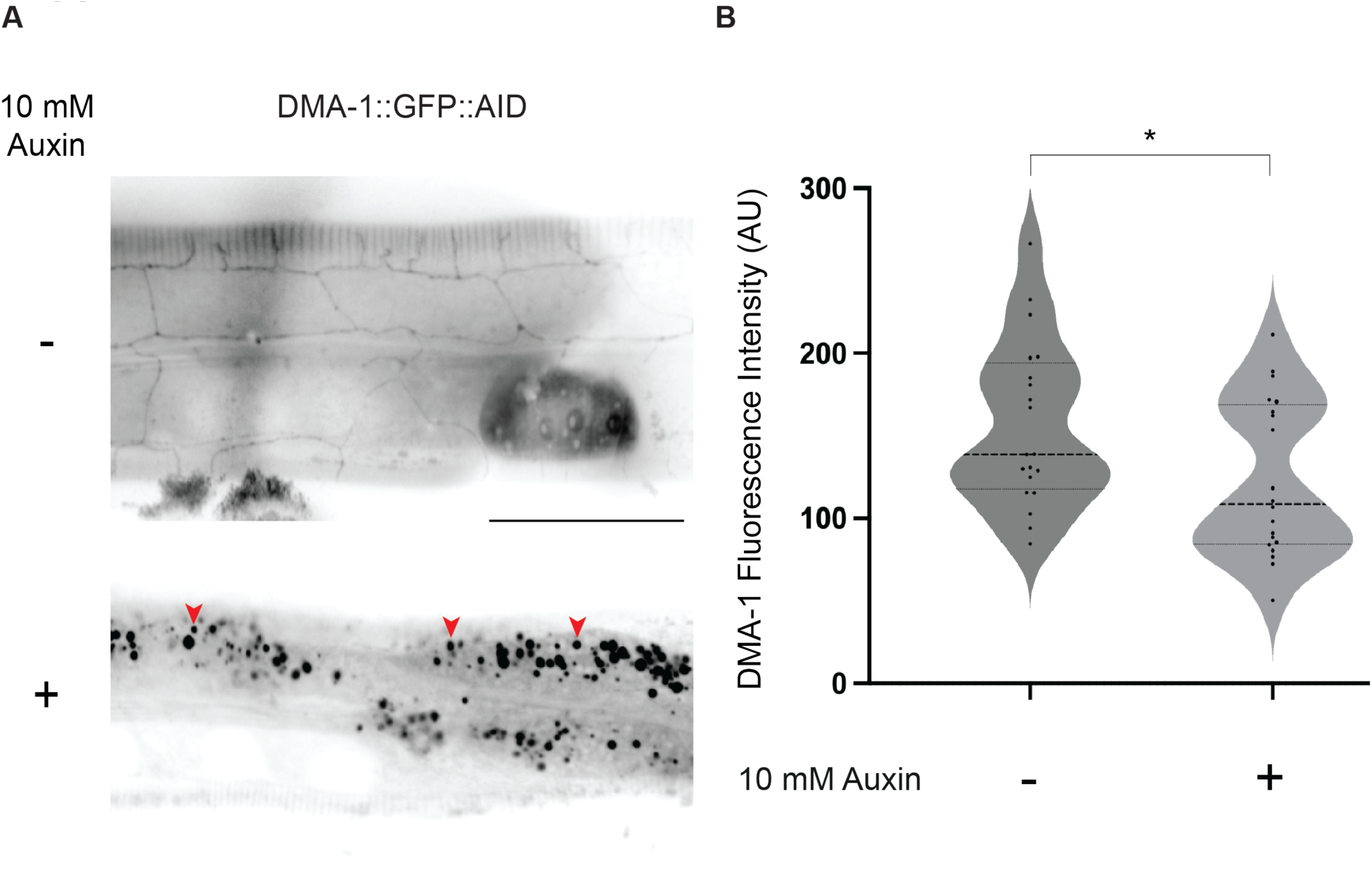
Temporal degradation of degron tagged DMA-1. (A) Lateral fluorescence sum intensity z-projections of endogenously labelled DMA-1::GFP::AID treated without auxin (top) or with 10 mM Auxin (bottom). Scale bar, 50 μm. Red arrowheads indicate examples of gut granules which exhibit autofluorescence and are not present in the PVD neuron (B) Quantifications of DMA-1::GFP::AID fluorescence intensity in the PVD cell body. Medians are represented in thick dashed lines and quartiles are represented in thin dashed lines. P value was calculated using a two-tailed unpaired Student’s *t*-test. n=20 for all conditions. *p≤0.05.

